# Uterine injury during diestrus leads to embryo spacing defects and perturbations in the COX pathway in subsequent pregnancies

**DOI:** 10.1101/2022.03.15.484521

**Authors:** Elisa T. Zhang, Kristen L. Wells, Lars Steinmetz, Julie C. Baker

## Abstract

Uterine injury from procedures such as Cesarean sections (C-sections) often have severe consequences on subsequent pregnancy outcomes, leading to disorders such as placenta previa, placenta accreta, and infertility. With rates of C-section at approximately 30% of deliveries in the US and that are projected to continue to climb, a deeper understanding of the mechanisms by which these pregnancy disorders arise and opportunities for intervention are needed. However, there are no animal models to date that comprehensively assess the consequences of uterine injury. Here we describe a rodent model of uterine injury on subsequent *in utero* outcomes. We observed three distinct phenotypes: increased rates of resorption and death, embryo spacing defects, and placenta accreta-like features of reduced decidua and expansion of invasive trophoblasts. We show that the appearance of embryo spacing defects depends entirely on the phase of estrous cycle at the time of injury. Using RNA-seq, we identified perturbations in the expression of components of the COX/prostaglandin pathway after recovery from injury, a pathway that has previously been demonstrated to play an important role in embryo spacing. Therefore, we demonstrate that uterine damage in this mouse model causes morphological and molecular changes, most notably perturbed expression of COX/prostaglandin pathway-related genes, that ultimately lead to placental and embryonic developmental defects.

## Introduction

Over the past decade, maternal mortality rates have risen in the United States. This escalation is due in part to placenta accreta spectrum, or PAS, in which the placenta invades too deeply into maternal tissues, even as far as the bladder^1,2^. Evidence suggests that PAS is linked to the condition placenta previa. In previa, the placenta is abnormally located in the lower portion of the uterus^3^. This placement of the placenta not only physically obstructs the cervix but is a significant risk factor for the more severe, invasive condition of PAS^2,4^. PAS and previa were rare prior to the 1930s but now occur in approximately 1 in 500 pregnancies^2,5–8^, likely due to the adoption of cesarean sections (C-sections) as a common practice^8–10^.

It is unknown how a C-section predisposes the uterine environment to previa and PAS, but some hallmarks of these diseases provide insight. One of the features of PAS is the absence of decidua between the invasive placental cells (trophoblasts) and the myometrium. This has led to the hypothesis that the decidua acts as a barrier, and if compromised, becomes permissive to extreme trophoblast invasion^4^. In both previa and PAS, increased numbers of invasive extravillous trophoblasts (EVTs) have been observed throughout the uterus, suggesting that the disease could manifest globally and not only at the site of injury^11–16^. As causal experiments are not possible in humans, animal models are an important tool to study the mechanisms underlying how uterine damage affects long-term maternal and fetal health. Studies in mice have shown that uterine injury, and subsequent wound healing, lead to changes in patterns of collagen deposition and biomechanical properties, including strength and elasticity, suggesting that localized damage could affect the overall function and contractility of the myometrium^17^. Importantly, studies have shown that uterine damage can lead to PAS features in subsequent pregnancies, including recent work from Burke et al. in 2020^18,19^. These reports strongly suggest that the mouse can be used to model how uterine damage subsequently predisposes pregnancies to previa and PAS in humans.

An important determinant of uterine biology is the extensive morphological and physiological changes that occur during the estrous cycle. The rodent estrous cycle repeats approximately every 4-5 days and is composed of four stages: proestrus, estrus, metestrus, and diestrus^20^. The uterus grows in thickness during proestrus in preparation for ovulation and mating receptivity during estrus and then shrinks as a result of endometrial degeneration during metestrus, resulting in a thin and elongated uterus at diestrus^21^. Reproductive hormones drive these anatomical transformations. Estrogen levels spike during proestrus and estrus and drop upon entering metestrus. Progesterone levels rise during metestrus, peaking during diestrus. Therefore, the uterus is a source of drastically changing hormones over the course of a 4-5 day period. While the estrous cycle is the rodent equivalent of the human menstrual cycle, with similar hormonal profiles that drive cyclical changes necessary for reproduction, there are some key differences, such as the lack of menstruation and pre-decidualization of the endometrium in rodents. Overall, the rapid hormonal fluctuations during the estrous cycle create dramatically different uterine morphologies that respond in distinct ways to the environment.

Embryo implantation in the mouse occurs with regular, systematic spacing between embryos along the two uterine horns. This regular spacing is defective in mice lacking components of the cyclooxygenase/prostaglandin (COX/prostaglandin) pathway, including *Lpar3*, *Pla2g4a*, and *Wnt5a*^*22–24*^. In mice homozygous mutant for each of these genes, embryos are clustered together, in some instances with shared decidua and fused placentas. In the COX/prostaglandin pathway, arachidonic acid is converted into bioactivate lipids known as prostaglandins via the activity of COX enzymes and specific terminal prostanoid synthases^25^ (Supplemental Fig. S4). Upon their biosynthesis, prostaglandins bind to specific prostanoid G protein-coupled receptors (GPCRs) to activate intracellular signaling and downstream gene transcription^25^ (Supplemental Fig. S4). Prostaglandins are known for their roles in inflammation, vasodilation, and reproduction^25–27^. Importantly, the major prostaglandin synthesized in the uterus (PGI_2_), as well as the uterine-specific PGF_2α_, stimulate smooth muscle contraction^28–31^. In addition, pharmacological inhibition of COX-2 in human myometrial tissue exerts significant relaxation in vitro^32^. Therefore, one hypothesis is that the COX/prostaglandin pathway regulates uterine contractility, which is necessary for the consistent spacing observed between mouse embryos.

In this study, we developed a mouse model to study mechanisms underlying previa and PAS. We injured mouse uteri by incision, examined future pregnancies, and found significant spacing defects, reduced embryo viability, and thinned decidua with increased numbers of trophoblast giant cells (TGCs). Importantly, we show that spacing defects occur up to 3 months post-injury and are entirely dependent on the mouse being in diestrus at the time of surgery, suggesting that there is a molecular memory of damage that is dependent on the estrous phase. To identify the contributors to this memory, we performed RNA-seq on uterine tissue post-surgery and found that components of the COX/prostraglandin pathway are impaired long-term after damage in diestrus, but not estrus. Overall, we find that defects in embryonic spacing are caused by uterine injury during diestrus and this leads to long-term misregulation of the COX/prostaglandin pathway.

## Results

### Novel mouse model of uterine injury

We developed a mouse model to study the effects of uterine damage on future pregnancies. To this end, we injured the left uterine horns of 7-12 week-old virgin females, leaving the right uterine horn intact (Fig. 1A; see Methods). In addition, we used two different controls: a sham surgery, in which mice underwent all procedural steps except the uterine cut, and no surgery, in which mice were not treated in any way. After a period of recovery (typically ~1 mo.), we bred the females and assessed the role of injury on placentation and fetal development. We observed distinct developmental defects during pregnancy as a result of previous uterine damage: increased rates of resorption and death, embryo spacing defects, and placentas with features of placenta accreta spectrum (PAS). Therefore, uterine damage affects subsequent pregnancies in mice in a multitude of ways that bear resemblance to human pregnancy conditions.

**Figure 1:**
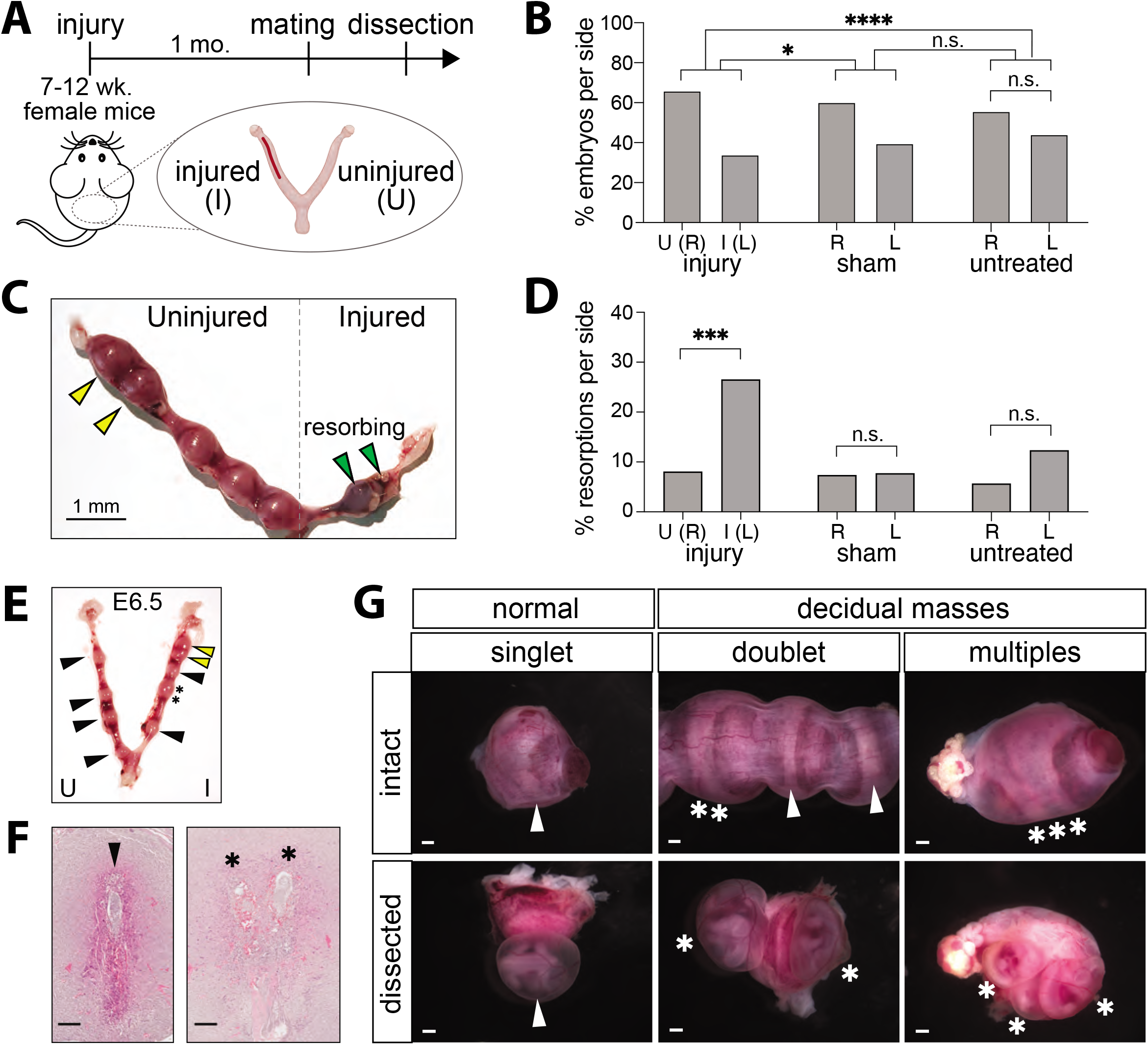
Uterine injury in mice leads to fewer implanting embryos, more resorptions, and embryo spacing defects. A. Schematic showing timeline of surgical procedure, mating and analysis. B. Percentage of total embryos in the injured (I or Left (L)) or uninjuried (U or Right (R)) uterine horn in animals subjected to surgeries (injury), to sham surgeries (sham), and to no treatment (untreated). (*) p < 0.05, (****) p < 0.0001. n.s. = not significant. C. E12.5 uterus showing the presence of resorbed embryos in the injured uterine horn. Scalebar = 1 mm. D. Percentage of resorbed embryos that are resorbed in the injured (I or L) or uninjured (U or R) horns in animals subjected to surgeries (injury), to sham surgeries (sham), and to no treatment (untreated). (***) p < 0.001. n.s. = not significant. E. Uterus showing the presence of misspaced embryos at E6.5. Embryos that have implanted as a decidual mass are indicated with stars. Other abnormally-spaced embryos are indicated with yellow arrows. F. H&E-stained sections of embryos from (E) showing a normal, single-embryo implantation site (left; embryo indicated by black arrow) and a decidual mass containing a doublet of embryos (right; embryos indicated by stars). Scalebar = 100 μm. G. Images showing intact, undissected uteri (top row) and dissected samples (bottom row) with normal singlets (arrows) or decidual masses containing a shared decidua capsularis (stars). Implantation sites contain either a single embryo (left), a doublet (middle), or triplet (right) are shown. Scalebar = 1 mm.

### Uterine damage inhibits fetal development in mice

To determine whether an injured uterus can support fetal development, we assessed the total number of embryos in the injured (left) and uninjured (right) horns. To this end, we dissected 63 pregnant mice at multiple stages of embryonic development (E12.5 (n = 39), E10.5 (n = 2), E11.5 (n = 6), E13.5 (n = 3), E14.5 (n = 2), E15.5 (n = 5), E16.5 (n = 1), E6.5 (n = 2), E8.5 (n = 3)). Across all 63 animals, we found a total of 179 (34%) embryos in the injured (left) horns and 342 (66%) in the uninjured (right) horns (p=1.27 x 10^−12^, one-sample proportions test compared to theoretical 0.5 proportion of embryos per horn; Fig. 1B). In the untreated no-surgery controls, we found 57 (44%) of the embryos on the left and 72 (56%) on the right. In sham surgery controls, we found 51 (40%) of the embryos on the left and 78 (60%) on the right. The reduced embryo numbers in the injured horns of the experimental condition is statistically significant in comparison to the observed embryo proportions in both the no-surgery (proportion of embryos in the left horn: 0.44; one-sample proportions test, p=1.13 x 10^−5^) and sham surgery controls (proportion of embryos in the left horn: 0.40; one-sample proportions test, p= 0.0098). In contrast, there is no difference in embryo numbers between the horns of sham surgery controls compared to untreated controls (p=0.35, one-sample proportions test) (Fig. 1B). In summary, we observed fewer embryos in the injured uterine horns.

The observation that fewer embryos develop in the injured horn may be due to either failure to implant or embryonic death. To determine whether the lower numbers were due to defects in implantation, we examined embryos at E6.5 and found no difference between the injured and uninjured horns (9 vs.10 embryos, injured vs. uninjured horns; p=1). We next examined the numbers of embryos at E8.5 and found fewer in the injured horn (9 vs. 21 embryos; p=0.045). All subsequent gestational ages had a similar bias, suggesting that injury does not affect the frequency of implantation, but increases the odds of embryonic death after E6.5. To understand rates of embryonic death, we next examined the health and viability of the developing embryos in each uterine horn. We found that 22.9% (n = 41/179) of embryos in the injured horns were in the process of resorbing, compared to 8.2% (28 / 342) of embryos in the uninjured horns (p = 4.86 x 10^−6^, two-sample test for equality of proportion) (Fig. 1C and 1D). In contrast, in sham surgery controls, the resorption rates were similar between the two horns, with 6.4% resorptions (n = 5/78) in the left horns and 7.8% (n= 4/51) in the right horns (p = 1, two-sample test for equality of proportion). Similarly, in the untreated no-surgery controls, the resorption rates were 7.7% (n = 4/52) on the left and 9.7% (n = 7/72) on the right. (p=0.30, two-sample test for equality of proportion) (Fig. 1D). These results show that there are fewer embryos in the injured horns and that many are undergoing resorption, suggesting that uterine injury frequently leads to fetal demise.

### Estrous phase at surgery dictates presence or absence of embryo spacing defects

In addition to fewer viable embryos developing in the injured horn, we observed spacing defects as early as E6.5 of gestation (Fig. 1E and 1F). By mid-gestation (E11.5 to E14.5), these presented as large decidual masses containing 2 and occasionally 3 embryos (Fig. 1G) and can be identified by a shared band of decidua capsularis encircling the yolk sac and amnion. Initially, we found only 50% of the injured dams contained these large decidual masses. Since the spacing defect after injury was not completely penetrant, we tested whether the estrous cycle was a factor. To this end, we performed surgery on mice during each phase of the estrous cycle - proestrus, estrus, metestrus, and diestrus - as determined by cytology of vaginal lavage^33^. We then examined the frequency of spacing phenotypes in pregnancy after surgery. We found that performing uterine surgery during estrus (estrus injury) led to no dams with spacing defects (n = 0/12) during pregnancy (Fig. 2A, 2B). However, we found that performing surgery during diestrus (diestrus injury) led to 90.1% (n = 20/22) of dams having spacing defects, as compared to 14.3% (n = 1/7) of sham surgeries performed during diestrus (Fig. 2A, 2B). Further, we found that surgery during proestrus and metestrus (proestrus- and metestrus-injury) led to 14.3% (n = 1/7) and 40% (n = 4/10) of dams having spacing defects, respectively (p= 9.69 x 10^−7^, diestrus vs. estrus; p = 0.0018, diestrus vs. metestrus; p = 0.00053, diestrus vs. proestrus; two-sample tests for equality of proportion) (Fig. 2A, 2B). Overall, this suggests that the diestrus phase predisposes the uterus to long-term effects of surgery.

**Figure 2:**
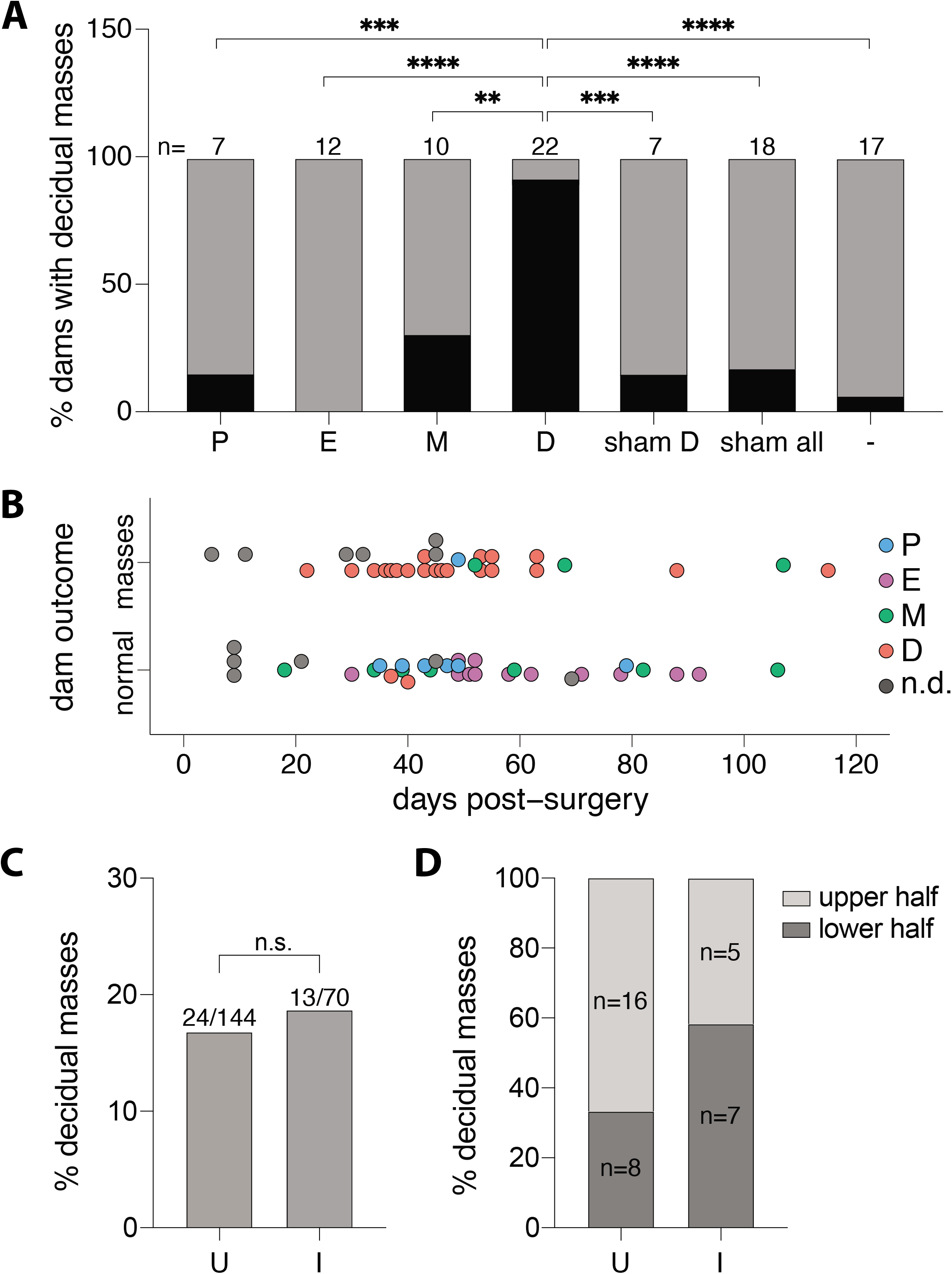
Embryo spacing defects are dependent upon phase of the estrous cycle at the time of injury. A. Percentage of dams displaying embryo spacing defects (black) after injury during proestrus (P), estrus (E), metestrus (M), or diestrus (D), or after sham surgery during diestrus (sham D), during all phases (sham all), or in untreated controls (−). B. Graph plotting effect of the estrous cycle at the time of surgery with the eventual appearance of decidua masses in pregnant dams from 0-120 days post-surgery. Number of days post-surgery is counted between date of surgery and date of plug (E0.5). Estrous phases are indicated as described in (A). C. Percentage of embryos contained in decidual masses in either the uninjured (U; left) or injured (I; right) horn. D. Percentage of decidual masses in the upper (light gray) compared to lower (dark gray) halves of each uterine horn (U = uninjured, I = injured).

### Embryo spacing is impaired regardless of time between injury and pregnancy

We next asked whether recovery time post-injury plays a role in subsequent embryo spacing defects. To this end, we analyzed spacing outcomes by length of recovery, which ranged between 5 - 107 days after uterine injury. We found that spacing defects occur regardless of recovery time and are observed even 107 days post-surgery (Fig. 2B). Importantly, in superimposing estrous phase onto recovery times, we again show that nearly all of the diestrus injuries, regardless of recovery time, resulted in embryo spacing defects, whereas estrus injuries never did. Overall, we find that recovery time has no discernible impact on outcome and that the estrous cycle is the only determinant of subsequent embryo spacing defects.

### Embryo spacing defects occur throughout the uterus

To determine whether the embryo spacing defects were specifically associated with a scarred uterus or whether they can arise anywhere within a damaged uterine environment, we assessed the distribution of the embryo spacing. To this end, we determined the percentage of implantation sites that presented as large decidual masses in both horns and found 18.6% (n = 13/70) in the injured horns and 16.7% (n = 24/144) in the uninjured horns (Fig. 2C). This indicates that spacing defects occur at the same frequency in both horns (p = 0.88, two-sample test for equality of proportion). We next examined the distribution of the large decidual masses within each horn. In the injured horns, we found 5/13 in the upper halves and 8/13 in the lower halves (Fig. 2D). Similarly, in the uninjured horns, we found 16/24 in the upper halves and 8/24 in the lower halves (Fig. 2D). Altogether, these results suggest that extreme spacing defects that result in large masses of decidua arise globally throughout the uterus.

### Embryo spacing defects affect placentation and embryogenesis

We next examined the embryos and placentas within these decidual masses. We found that placentas in shared concepti either remained separate (n = 20/37, Fig. 3A, left) or were fused together (n = 17/37; Fig. 3A, right). In both scenarios, by histology we detected extensive labyrinth defects with malformed blood sinuses within the smaller of these placentas (Fig. 3B-C). Further, we saw that most of the embryos within the decidual masses were underdeveloped, ranging from slightly smaller to fully resorbed with little or no embryonic tissue present (Fig. 3D). We investigated the extent of these growth defects by measuring the weights of embryos at mid-gestation (E11.5 – E14.5) (Fig. 3E, Supplemental Fig. S1A-C). We found that the embryos within the decidual masses weighed less than their littermates (1.052 vs. 0.881, mean singleton vs. misspaced embryo weights, normalized to mean weight per litter; p = 0.0007, two-tailed t-test) (Fig. 3E). In addition to decreased embryo weight, we also observed an instance in which a misspaced embryo developed a twisted body axis *in utero*, presumably as a result of mechanical constraints in a crowded environment (Supplemental Fig. S1D). Altogether, we demonstrate that the decidual masses that form as a result of uterine injury are associated with both placental and embryonic insufficiency.

**Figure 3:**
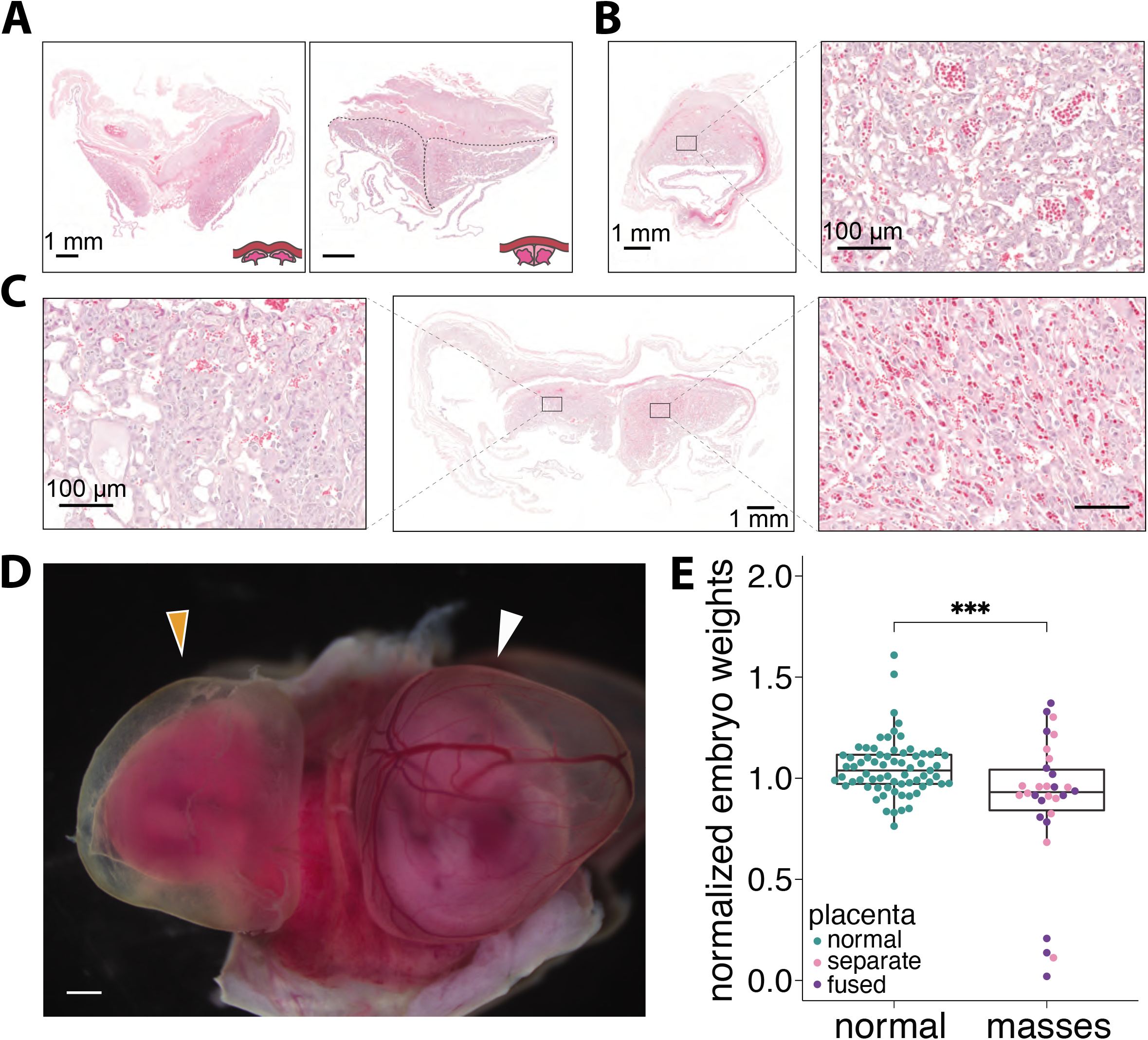
Embryo spacing defects affect placentation and embryogenesis. A. H&E-stained sections of decidual masses containing separate (left) and fused (right; dashed gray lines) placentas. B. H&E-stained section of a normal placenta with uterus attached. A magnified view of the placental labyrinth is shown in the inset. Scalebar = 1 mm; inset scalebar = 100 μm. C. H&E-stained section of placentas in a decidual mass showing labyrinth defects in the placenta on the left. Scalebar = 1 mm; inset scalebar = 100 μm. D. Images of embryos within a decidual mass containing one underdeveloped embryo (left, orange arrow) and one normal embryo (right, white arrow). Scalebar = 1 mm. E. Weights of embryos collected at E11.5-E14.5 normalized to mean litter embryo weight. Embryos not in decidua masses (normal; green dots) compared to embryos within decidual masses (masses; either pink or purple dots). Pink dots represent placentas in decidual masses that remain separate. Purple dots represent placentas in decidual masses that are fused (p=0.0007).

### Placental fusion requires a male genotype

Since the decidual masses contained multiple embryos, we next asked how they arose. Two possibilities could explain this phenomenon: either separate embryos implanted very close together, or a single embryo twinned. To distinguish between these scenarios, we determined the sexes of 36 embryo pairs and performed PCR for the sex chromosomes (see Methods, Supplemental Fig. S2). We detected a total of 15 pairs that contained both a male and a female, suggesting that two separate embryos implanted very closely together to give rise to these decidual masses (Supplemental Fig. S2A). Of the 36 embryo pairs, 17 were associated with placentas that were fused and 19 with separate placentas. When we examined the sexes of the 17 embryo pairs with fused placentas, (Fig. 3A, Supplemental Fig. S2) we found that 8 were male-male, 9 were male-female, and none were female-female (Supplemental Fig. S2, p = 0.02, Chi-square test). In contrast, when we examined the sexes of the 19 pairs with separate placentas, we found that 6 were male-male, 6 were male-female, and 7 were female-female (Supplemental Fig. S2, p = 0.26, Chi-square test). Taken together, these results indicate that embryos within decidual masses are the result of severe spacing defects of separate and not twin embryos at implantation. Further, we show that placental fusion resulting from uterine injury occurs only in the presence of at least one male embryo.

### Uterine injury leads to reduced decidua and increase in TGCs

We observed that placentas from injured animals were difficult to separate from the uterine wall during dissection, suggesting an abnormal attachment between the maternal and fetal tissues. To understand the cause of this adherence, we histologically examined 81 individual placental-uterine attachments from both injured and uninjured horns from 25 pregnant mice. We found that 14 of the 81 placental-uterine units had a reduced decidual layer, of which 7 had no decidual stromal cells and only the mesometrial lymphoid aggregate of pregnancy (MLAp) (Fig. 4A, B, and C). 9 of these 14 were found in injured horns, whereas 5 were in uninjured horns. In comparison, we did not observe this phenotype in any of the untreated no-surgery controls (n = 0/26; p = 0.05, two-sample test for equality of proportion). We quantified decidual thickness by measuring the distance at the midpoint of the placenta between the top of the trophoblast giant cell (TGC) layer (or the top of spongiotrophoblast (SpT) layer, in instances with no TGCs present) to the bottom of the myometrium and observed a statistically-significant difference between the 14 from injured animals and the 26 from no-surgery controls (p = 2.87 x 10^−14^, Welch’s t-test) (Fig. 4C). In histological sections of these 14 maternal-fetal interfaces, we also observed more TGCs (Fig. 4B), which are highly polyploid cells that contain large nuclei identifiable by H&E staining. To quantify this observation, we counted the number of TGCs in the middle third of placentas, where few TGCs remain by mid-gestation during normal development (Fig. 4A and B, insets). We found nearly ten times the number of TGCs compared to those from untreated controls (mean of 12.4 TGCs in placentas associated with reduced decidua, compared to 1.7 TGCs in placentas from untreated controls, p = 3.01 x 10^−5^, Welch’s t-test) (Fig. 4D). Therefore, in a subset of placentas from injured animals, we find evidence of both reduced decidua and expanded numbers of TGCs, features that resemble two key hallmarks of PAS in human patients.

**Figure 4:**
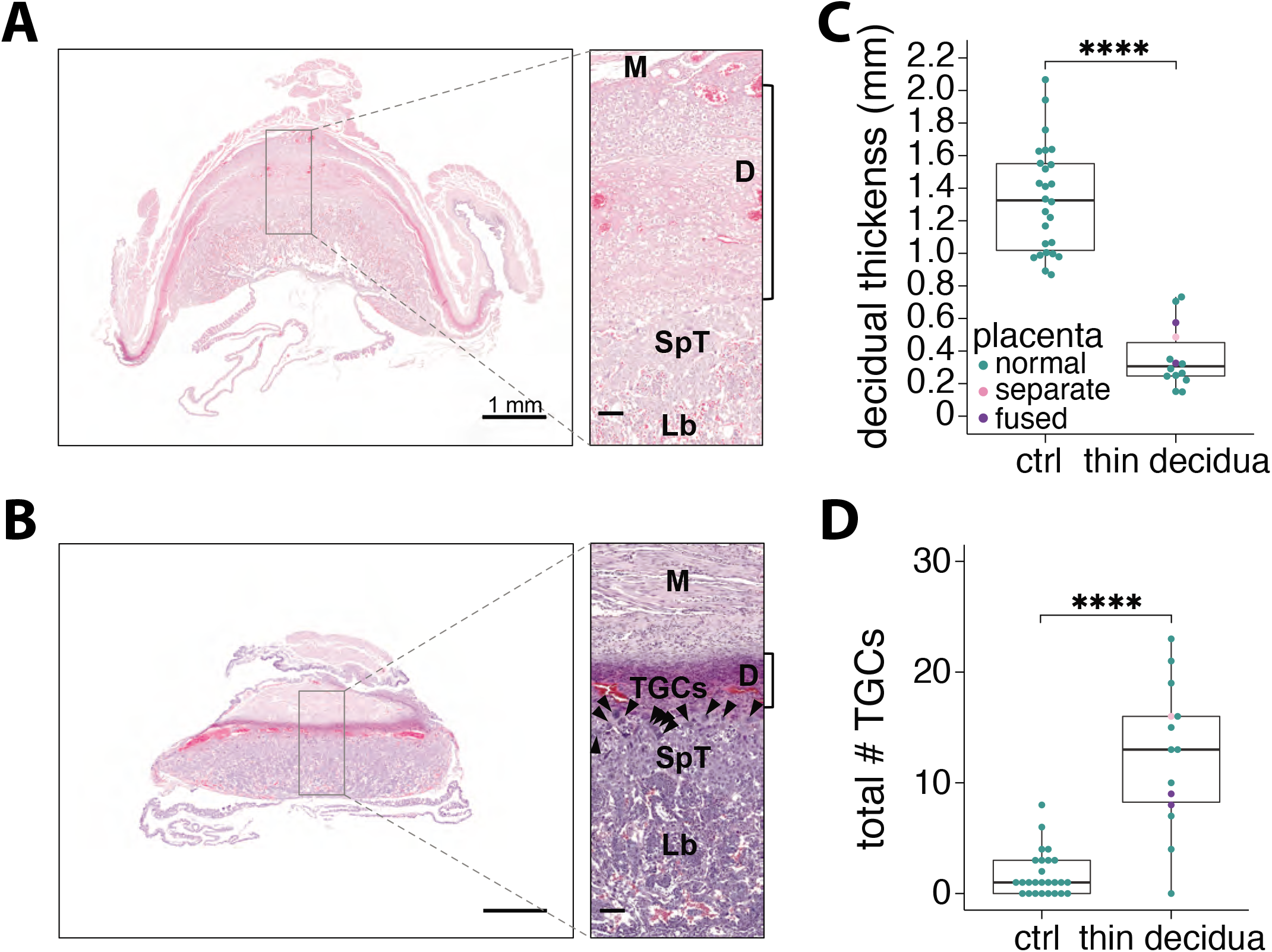
Uterine injury leads to placentas with features of PAS. A. H&E-stained section of a normal placenta attached to the uterus. Myometrium (M), decidua (D), spongiotrophoblast (SpT), and labyrinth (Lb) are indicated. Scalebar = 1 mm; inset scalebar = 100 μm. B. H&E-stained section of a placenta with reduced decidual thickness and expanded numbers of trophoblast giant cells (TGCs). Scalebar = 1 mm; inset scalebar = 100 μm. C. Decidual thicknesses of normal placentas from control dams (ctrl) and affected placentas from injured dams. (****) p < 0.0001. D. Total number of TGCs in normal placentas from control dams (ctrl) and affected placentas from injured dams. (****) p < 0.0001.

### Injuries performed in diestrus heal less efficiently

We next sought to determine how the uterus heals after diestrus and estrus injuries (Fig. 5). We macroscopically and histologically examined wounds 3, 5, 7 days or 1 month after injuries (n=4-5 per timepoint per condition) (Fig. 5). By day 3, wounds from estrus injuries had closed (Fig. 5, top row). In contrast, by 3 days after diestrus injuries, wounds remained open and were often accompanied by distension, particularly of the endometrial layer, and inversion (Fig. 5, top row). Inflammation with infiltrating lymphocytes, as well as partial or complete absence of the luminal epithelium at the wound, were seen after both diestrus and estrus injuries (Fig. 5, top row). By day 5, the wounds from diestrus injuries had closed (Fig. 5, second row). The wounds from both diestrus and estrus injuries contained a continuous endometrial layer, nearly-apposed inner (circular) and outer (longitudinal) myometrial layers, and inflammation with infiltrating lymphocytes, as well as scar-like tissue with residual edema on the outer serosal surface (Fig. 5, second row). However, the scars from estrus injuries were smaller than those from diestrus injuries. By day 7, the wounds from estrus injuries were shallower than those at day 5 but still smaller than those from diestrus injuries (Fig. 5, third row). By 1 month, the wounds from diestrus injuries were similar to those from estrus injuries (Fig. 5, bottom row). Hemosiderin is present in both conditions (Fig. 5, bottom row), as are lymphocytes, though these are fewer in number compared to earlier timepoints (Fig. 5). While extracellular matrix changes, misaligned muscle fibers, and fibrosis can be found 1 month post-injury, the endometrial and myometrial layers have closed fully, suggesting that the uterus contains remarkable wound healing capabilities that can largely restore the overall structure of the uterus with no intervention (Fig. 5, bottom row). Taken together, these experiments identify differences in wound healing between diestrus- and estrus-injured animals, particularly in the days immediately following injury.

**Figure 5:**
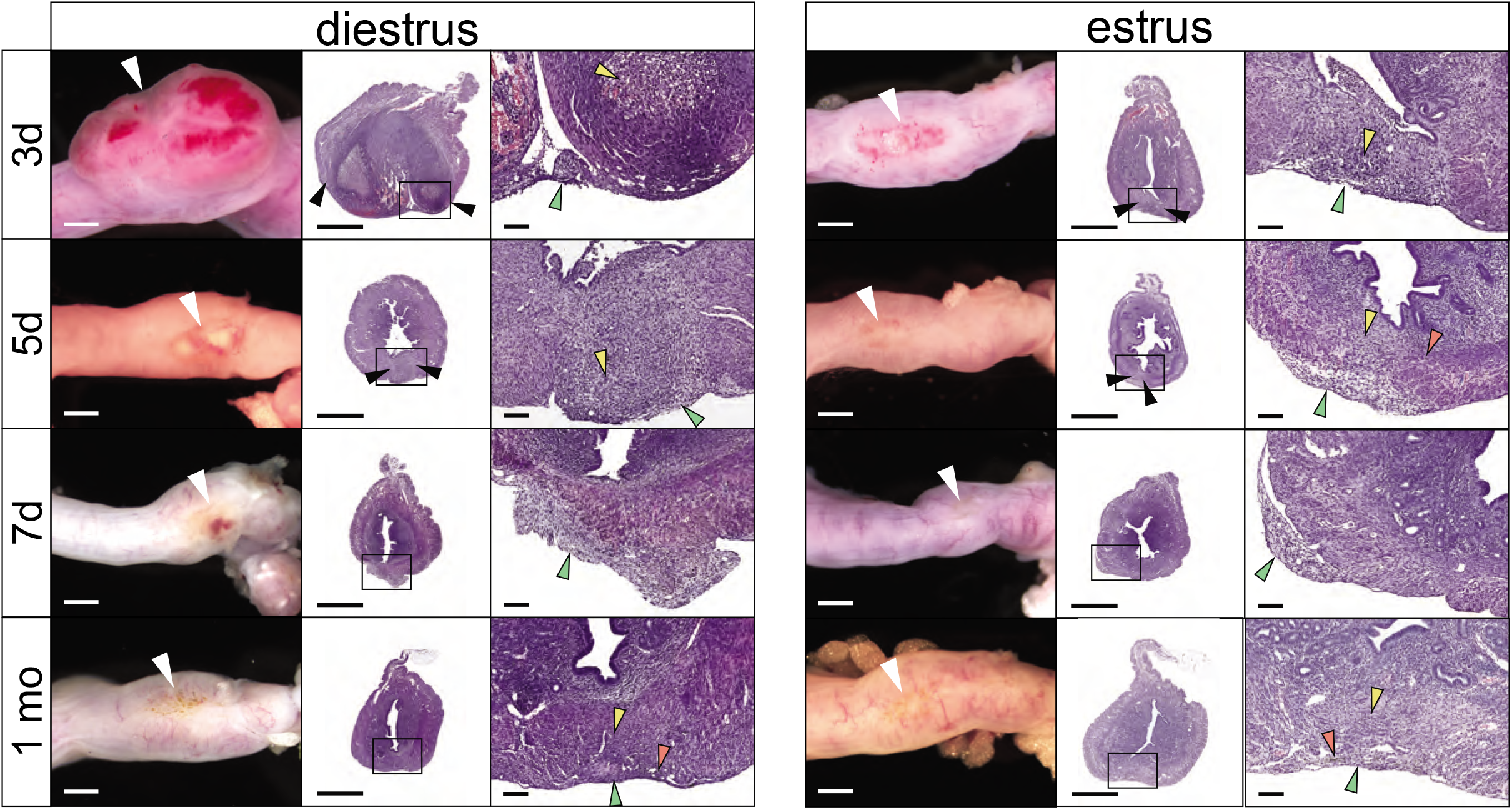
Uterine wound healing properties depend on estrous phase at time of injury. Images (left) and corresponding H&E-stained sections (right) of uteri injured during either diestrus or estrus, with time of recovery indicated on the far left. White arrows indicate scar site. Black arrows indicate myometrial borders at the sites of prior incision. Green arrows indicate scar tissue. Yellow arrows indicate areas of inflammation with lymphocytes, and red arrows indicate hemosiderin. Scalebar = 1 mm; inset scalebar = 100 μm.

### The COX/prostaglandin pathway is misregulated in diestrus-injured horns

We next investigated whether there was a molecular signature caused specifically by uterine injury in diestrus. To this end, we isolated uterine mid-sections from mice that had been injured during diestrus (n = 3), estrus (n = 3), as well as no surgery controls (n = 4). We then performed RNA-seq on the injured (left) and uninjured (right) horns for each experimental animal and both horns from the controls. To determine the overall relationship between these samples, we first performed principle component analysis (PCA). This revealed that each diestrus-injured horn displayed a systematic shift along the first principle component (PC1) relative to the uninjured horn from the same animal (Fig. 6A), whereas estrus-injured horns clustered more closely to their corresponding uninjured horns. However, we also observed strong individual-specific effects, as uterine horns from the same animal showed the strongest similarities (Fig. 6A, gray ovals). After correcting for individual variation, the PCA again shows that the injured and uninjured horns are more distinct from each other following surgeries during diestrus compared to estrus (Fig. 6B). The horns from the no-surgery controls were indistinguishable from one another by PCA (Fig. 6A and B; n = 2). This suggests that injury during diestrus establishes a more distinct and lasting molecular signature of damage compared to estrus (Figs. 5 and 6).

**Figure 6:**
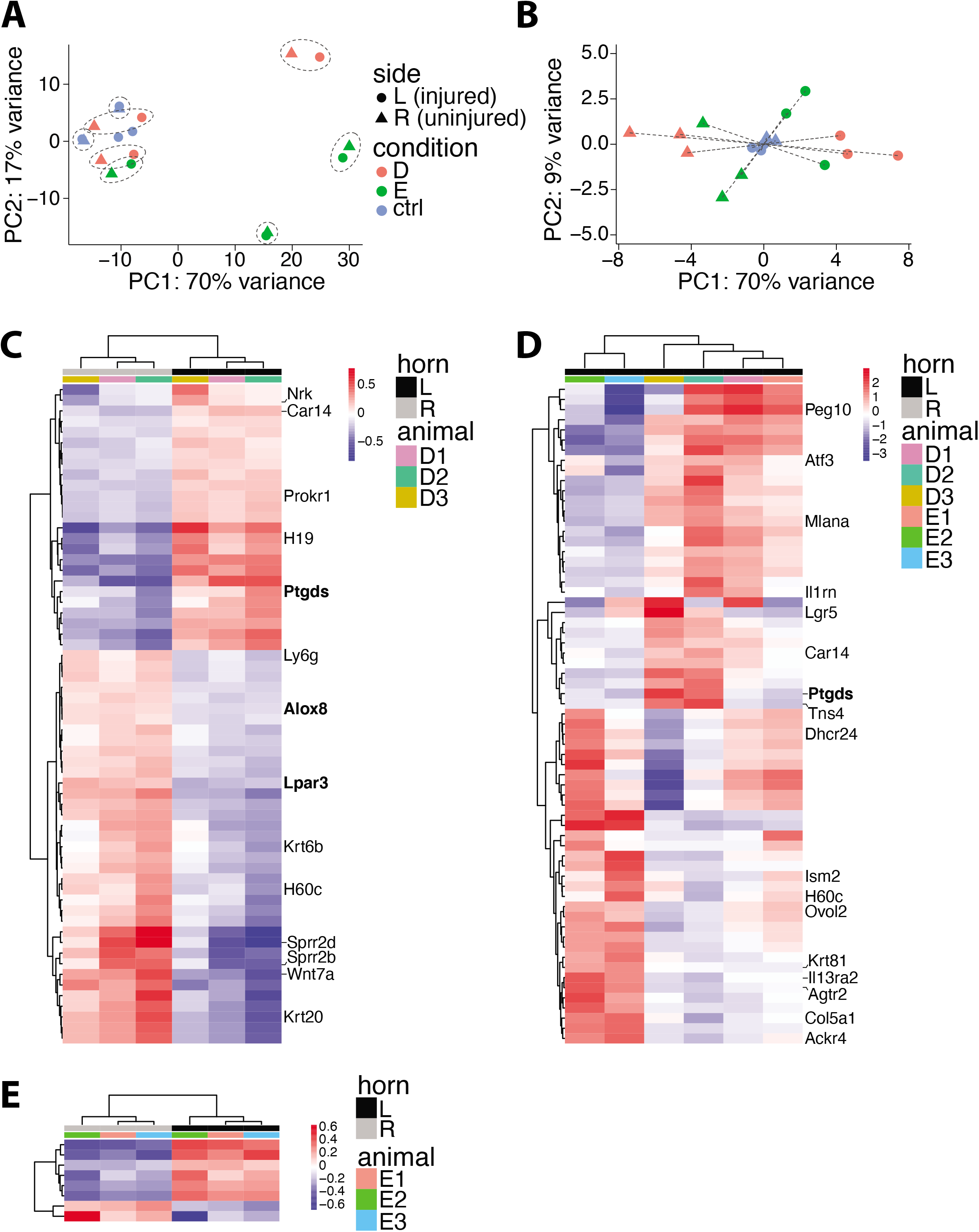
RNA-seq reveals molecular signatures of diestrus injury. A. Principle component analysis (PCA) plot of RNA-seq samples. Samples from the two uterine horns of the same dam are indicated with dotted gray lines. The injured (L) uterine horn of each dam is indicated by a triangle; the uninjured (R) uterine horn is indicated by a circle. Dams in diestrus (D) or estrus (E) at time of surgery are indicated in red and green, respectively. Control dams are colored in blue. B. PCA plot of samples after correcting for individual variation. C. Heat map showing differentially-expressed genes (DEGs) between the diestrus-injured and -uninjured uterine horns with a fold-change > 2. COX/prostaglandin pathway genes are shown in bold. D. Heat map showing differentially-expressed genes (DEGs) between the diestrus-injured and estrus-injured uterine horns with a fold-change > 2. COX/prostaglandin pathway genes are shown in bold. E. Heat map showing differentially-expressed genes (DEGs) between the estrus-injured and -uninjured uterine horns with a fold-change > 2.

We identified 62 transcripts that are misexpressed in the uterus following diestrus injury after correcting for individual variation (DEGs; 25 upregulated and 37 downregulated; adjusted p <0.05 and a >2-fold change) (Fig. 6C). We found that *Ptgds*, *Lpar3, and Alox8*, which are components of the COX/prostaglandin pathway, were differentially expressed between the diestrus-injured compared to diestrus-uninjured horns (Fig. 6C, Supplemental Table S2 for complete list; Supplemental Fig. S3B). We then compared diestrus-injured to estrus-injured horns. We found 65 DEGs (32 upregulated and 33 downregulated) that exhibited >2-fold change (adjusted p < 0.05) (Fig. 6D, Supplemental Table S3 for complete list). Among the 32 upregulated genes, we again identified *Ptgds*. Since previous studies have implicated COX/prostaglandin pathway as an important regulator of embryo spacing^22–24^, the presence of these components amongst our DEGs points to a molecular mechanism by which embryo spacing defects may arise.

We next compared estrus-injured and -uninjured horns and found only 8 DEGs (6 upregulated and 2 downregulated; adjusted p <0.05, >2-fold change), which is consistent with their overall similarity seen by PCA (Fig. 6A, 6B, and 6E). This smaller number of DEGs further highlights the minimal impact of estrus injury on uterine tissue (Fig. 5C, Fig. 6A and B). The absence of extensive differences between estrus-injured and -uninjured horns is also observed when we only use an adjusted p-value <0.05 with no fold-change threshold. In this scenario, only 57 DEGs are found (Supplemental Fig. S3A). In contrast, when we perform this same, less stringent analysis between the diestrus-injured and -uninjured horns (adjusted p <0.05 with no fold-change threshold), we find 2084 DEGs (Supplemental Fig. S3B). Overall, this supports our finding that there is minimal long-term impact, either morphologically, functionally, or molecularly, as a consequence of estrus injury. In contrast, there are molecular charges that occur upon diestrus injuries, particularly within the COX/prostaglandin pathway, which has previously been implicated in embryo spacing.

## Methods

### Mouse injury surgical procedure

All mice were housed under a 12:12 light-dark cycle. All mouse procedures were performed in accordance with the APLAC protocol and institutional guidelines set by the Veterinary Service Center at Stanford University. All mice were obtained from Charles River or Jackson Laboratories. 100 mg/kg of ketamine and 10 mg/kg of xylazine were delivered intraperitoneally as anesthesia. After shaving the incision site, cuts were made through the skin and the fascia to expose the uterus. Injury was induced by inserting a burred 25G needle just beneath the uterotubal ligation into the lumen of the uterus and scraping the interior of the uterus along the anti-mesometrial surface until a full-thickness cut was generated. No major intra- or post-operative complications were observed in this study.

### Determination of estrous stage

Estrous cycle determinations were conducted by cytology of vaginal lavage, as adapted from the procedures described in McLean et al. 2012^33^. Lavage samples (~50 uL of sterile water) were spotted on glass slides, allowed to air-dry at room temperature, and then staining for 1 minute with a 0.1% crystal violet solution in 2% EtOH, followed by a 1-minute wash in distilled water and subsequent mounting with a coverslip.

### Timed matings

Copulation was determined by the presence of a vaginal plug the morning after mating, and embryonic day 0.5 (E0.5) was defined as noon of that day.

### Determination of embryo sex

Genomic DNA from ~5-10 mg of embryonic was extracted in 200 uL of PBND buffer (50 mM KCl, 10 mM Tris pH 8.3, 2.5 mM MgCl_2_, 0.1 mg/mL gelatin, 0.45% v/v NP-40, 0.45% v/v Tween-20) + 10 μg of proteinase K overnight at 55°C. Genomic DNA from 4-8 microdissected 10 μm FFPE sections were extracted using the QIAamp DNA FFPE Tissue kit (Qiagen, cat. no. 56404) according to manufacturer instructions. Sex was determined by PCR using the forward primer 5’-CCGCTGCCAAATTCTTTGG-3’ and the reverse primer 5’-TGAAGCTTTTGGCTTTGAG-3’ to detect the presence of the Smcx gene on the X chromosome (330 bp) in females and males, in addition to the Smcy gene on the Y chromosome (290 bp) in males.

### RNA-seq

~2-3 mm long uterine sections collected from the midpoint of the uterine horn encompassing all tissue types (myometrium and endometrium) were placed into TRIzol Reagent (Life Technologies, Cat. #15596) and stored at −80 degrees until extraction. Tissue homogenization was assisted with a motorized pestle mixer (Argos Technologies, cat. # A0001). RNA was then isolated according to the manufacturer’s instructions (Life Technologies, Cat. # 15596). RNA was quantified using a Nanodrop 1000 (Thermo Scientific) and the Qubit RNA High-Sensitivity (HS) Assay kit (Invitrogen, Cat. # Q32852). RNA quality was assessed using a Bioanalyzer 2100 (Agilent), and samples with an RNA Integrity Number (RIN) of 7.0 or greater were used for RNA-seq. Strand-specific, poly(A)-selected RNA-seq libraries were prepared by Novogene Co. using the NEBNext UltraII RNA library prep kit for Illumina (cat. #E7775) and sequenced on a NovaSeq6000 (Illumina) to produce paired-end 150-bp reads (average 40.4 million reads per sample). RNA-seq reads were mapped to the mm10 (GRCm38.p4) reference genome (Supplemental Table S1) using STAR version 2.6.1d^34^ to quantify gene expression. Read counts were quantified using featureCounts^35^ from the subread package (version 1.6.4), which identified a total of 27,999 transcripts. Transcripts with greater than 10 reads across all samples were included for further analysis. Differential gene expression analysis was performed using DESeq2 (version 1.30.0)^36^. To account for animal-specific batch effects, the following design was used when creating the DESeq object formula (~horn + horn:animal + horn:treatment), where horn refers to the left or right uterine horn, animal refers to the specific mouse, and treatment refers to whether the horn was injured or uninjured. After object construction with this formula, differential expression was performed using an adjusted p-value cutoff of 0.05 and a log_2_ fold change cutoff of 1.

### Histological analysis

Uterine and placental samples were collected by surgical removal, fixed overnight in 4% PFA (EMS, cat. #), dehydrated in a graded series of ethanol solutions, embedded in paraffin (Leica Surgipath Paraplast, cat. # 39601006), sectioned at 5 μm thickness, mounted on Superfrost glass slides (Fisher Scientific, Cat. #12-550-15), and stained with hematoxylin and eosin (H&E) using standard procedures of deparaffinization, rehydration via a graded series of ethanol solutions, stained with Harris hematoxylin (Sigma-Aldrich cat. # SLBV6928), dehydrated to 70% ethanol, stained with eosin (Sigma-Aldrich cat. # SLBW3770), fully dehydrated via a graded ethanol solution series to 100% ethanol, followed by mounting with either Acrymount (StatLab cat. # SL80-XX) or Cytoseal 60 (ThermoFisher Scientific cat. # 8310). Slides were imaged using either a Philips IntelliSite Ultra Fast or a Leica AT2 digital slide scanner.

### Statistical analysis and graphics programs

Statistical tests in this study were performed in either R v.3.5.2 or GraphPad Prism. Graphical illustrations were created with the help of BioRender.com.

## Discussion

Defective embryo spacing was the most salient phenotype we observed following uterine damage. These spacing defects led to fused placentas and high rates of embryonic death. Despite the injury being confined to the left horn, embryo spacing defects were identified in both the injured and uninjured horns, suggesting that localized molecular changes at the site of injury has global consequences throughout the entire uterus. Since the COX/prostaglandin pathway controls uterine contractility, which in turn regulates embryo spacing, we postulate that perturbations in the COX/prostaglandin pathway following localized damage thus translates into global contractility defects that impact the entire uterine environment. Intriguingly, embryo spacing defects were highly dependent on the phase of the estrous cycle during which injury was induced, suggesting that differences in uterine properties across the estrous cycle play a role in dictating both uterine wound healing and subsequent pregnancy outcomes. While diestrus-injured animals were nearly universally susceptible to such defects, estrus-injured animals were unaffected. This bias was not dependent upon time post-surgery, and indeed animals bred out >100 days still displayed the phenotype, strongly suggesting that the uterus retains a “memory” of damage incurred during diestrus but not estrus. This is particularly striking given that few structural differences remain between diestrus- and estrus-injured uteri after 1 month of recovery. Overall, this suggests that initial events post-injury in the uterus leave a global molecular “memory” that subsequently determines embryo spacing.

In searching for the molecular basis of this memory, we found that misregulation of the COX/prostaglandin pathway may underlie the long-term consequences of uterine damage. Of the 62 DEGs that met a significance threshold of an adjusted p < 0.05 and a fold-change threshold >2, we found the COX/prostaglandin pathway members *Lpar3* and *Alox8* to be downregulated after diestrus injury, while *Ptgds* (Prostaglandin D2 Synthase) was upregulated.

When we expanded our view to include all 2084 DEGs of any fold-change that reached an adjusted p < 0.05 threshold, we identified an additional 7 COX pathway components amongst the DEGs: *Ptgs2* (COX-2), *Pla2g4a*, *Ptges2* (Prostaglandin E Synthase 2), *Ptgfrn* (Prostaglandin F2 Receptor Inhibitor), *Alox5*, *Wnt5a*, *Ptger3* (Prostaglandin E Receptor 3 or EP3), and *Ptgis* (Prostaglandin I2 Synthase). The preponderance of members of this biosynthetic pathway specifically in the diestrus-injured horn suggests a role in the embryo spacing phenotype. Importantly, this is strongly supported by genetic and pharmacological experiments demonstrating that the prostaglandin biosynthetic pathway regulates embryo spacing. For example, administration of the prostaglandin inhibitor indomethacin during implantation impairs embryo spacing in rats^37^. Deletion of cytosolic phospholipase A2alpha (cPLA2α), which generates arachidonic acid (AA) for the COX enzymes, results in aberrant embryo spacing^24^. Similarly, upstream regulators of the COX/prostaglandin pathway such as *Lpar3* (LPA3)^23^, *Wnt5a*^22^, and adrenomedullin (*Adm*)^38–40^ also lead to defective embryo spacing upon knockout. (Supplemental Fig. S4). Overall, the link between the COX/prostaglandin pathway and the regulation of embryonic spacing, coupled with the identification of many of these pathway molecules specifically in the diestrus-injured horns, implicates COX/prostaglandin signaling as the causal uterine defect that is established after damage and that leads to long term issues during pregnancy.

It has been hypothesized that the COX/prostaglandin pathway regulates uterine contractility to facilitate proper embryo spacing in the uterus. The role of prostaglandins in regulating smooth muscle contractions is well documented. Lysophosphatidic acids (LPAs) signal through receptors such as LPA3 to induce smooth muscle contraction^41,42^, including the smooth muscle of the uterus^43^. Importantly, the major prostaglandins synthesized in the uterus, PGI_2_ and PGF_2α_, stimulate smooth muscle contraction^28–31^. In addition, the LPA3-selective agonist T13^44^ induces contractile responses in WT but not *Lpar3*^*−/−*^ mouse uteri^45^, and inhibition of COX-2 in human myometrial tissue results in substantial relaxation *in vitro* (Supplemental Fig. S4). Therefore, previous studies have implicated COX pathway genes and their control of uterine contractility as important regulators of embryo spacing. The question remains how this pathway is perturbed after diestrus injury. We suggest that diestrus injury irreversibly alters a key upstream regulator, causing persistent errors throughout the biosynthetic pathway through epigenetic or other mechanisms, thus altering contractility of the uterus and eventually leading to embryo spacing defects.

We found that diestrus and estrus uteri had different wound healing trajectories, with diestrus-injured uteri being substantially slower to heal than estrus-injured uteri (Fig. 5) in the first 7 days after damage. Several mechanisms could account for this slow initial healing. First, since estrus uteri are much thicker than diestrus uteri, the two cut surfaces of the uterus may be better able to contact each other to facilitate faster healing^46^. Second, estrus uteri display more frequent uterine contractions, which have been shown to promote endometrial healing^47^. Third, higher rates of cell proliferation in the estrus uterus may facilitate faster wound healing. Fourth, ECM, immune, and/or cell signaling genes expressed during diestrus and estrus could also drive wound healing properties. Lastly, estrogen and progesterone, which are at higher levels during estrus and diestrus, respectively, may have profound contributions. Studies have shown that while estrogen may enhance wound healing^48,49^, progesterone is inhibitory^50–53^. This is consistent with the open wounds we observed 3 days post-injury in diestrus, a phase characterized by a high progesterone:estrogen ratio^20,21^. Despite the initial slow healing in diestrus, the uterus completely heals by one month regardless of estrous phase at the time of injury. This is consistent with other rodent studies demonstrating complete closure of wounds within 6-8 days^46^ and the absence of histological scars between 14-25 days^46^ or at 60 days after injury^17^. Taken together, slower-healing wounds following diestrus injuries may arise from a variety of mechanical and cellular factors.

One of the consequences of improper embryonic spacing is the fusion of the placenta, which appears to require a Y chromosome. In all instances of placental fusion, the placentas were either male:male or male:female. We did not find female:female fusions. Intriguingly, there is an association between human previa and male sex (1.19 male:female ratio in women with previa vs. 1.05 in women without, p<0.001)^54^, suggesting that maleness creates an environment permissive for incorrect uterine spacing. This is unlikely to be due to testosterone since testosterone production in male embryos does not begin until mid-gestation, peaking only at late gestation^55–58^. However, our finding does imply that male-specific factors are involved. In addition to uterine factors, embryo-specific factors have been shown to influence spacing as transferring BPH/5 embryos into BPH/5 mothers, but not C57BL/6 embryos into either BPH/5 or C57BL6 mothers, nor BP5/H embryos into C57BL/6 mothers, caused spacing defects^59^, suggesting that forming the correct embryo:uterine attachment is highly specialized. Since it is known that placentas have sex-specific differences, it will be interesting to pursue those that bias uterine spacing defects. A consequence of these placental fusions include smaller placentas and embryos, as well as branching defects in the placental labyrinth layer. Overall, these findings show that embryo spacing defects have consequences on placental and fetal development and that a number of factors, including maleness, may contribute to differences in severity of the spacing problems.

While mouse physiology is clearly distinct from that of humans, the phenotypes arising after uterine damage in diestrus have several parallels to human previa and PAS. In previa, the placenta develops in the lower as opposed to upper segment of the uterus. Importantly, previa arises as a result of prior uterine damage^2,10^, suggesting that uterine injury may recapitulate common mechanisms giving rise to both misspacing in mice and previa in human. Further, in this mouse model of uterine injury, placentas have thin decidua and increased numbers of invasive TGCs, which bear obvious phenotypic parallels to human PAS. This mouse model of uterine injury could also be used to study vanishing twin syndrome in humans, whereby two embryos implant too closely together, resulting in the death of one twin and, on occasion, adverse outcomes for the surviving twin. In these scenarios, we suggest that investigation into the involvement of the COX/prostaglandin pathway, as well as estrogen and progesterone, would be warranted. While there has been no study to date to determine the impact of menstrual cycle phase on non-gravid uterine procedures such as myomectomy, the timing of C-sections has been shown to impact subsequent pregnancy outcomes. C-sections performed prepartum but not intrapartum led to elevated risk of previa and accreta^30^, suggesting that the state of the uterus at the time of injury may impact its healing and subsequent function. Importantly, the impact of estrous and menstrual cycles on uterine biology and uterine wound healing is an important avenue for further investigation, and mouse injury models may enable a mechanistic examination of human conditions associated with uterine injury, including previa, vanishing twin syndrome, and PAS.

