## Supplemental Fig. S1 for "Uterine injury during diestrus leads to embryo spacing defects and perturbations in the COX pathway in subsequent pregnancies"

Supplemental Figure S1

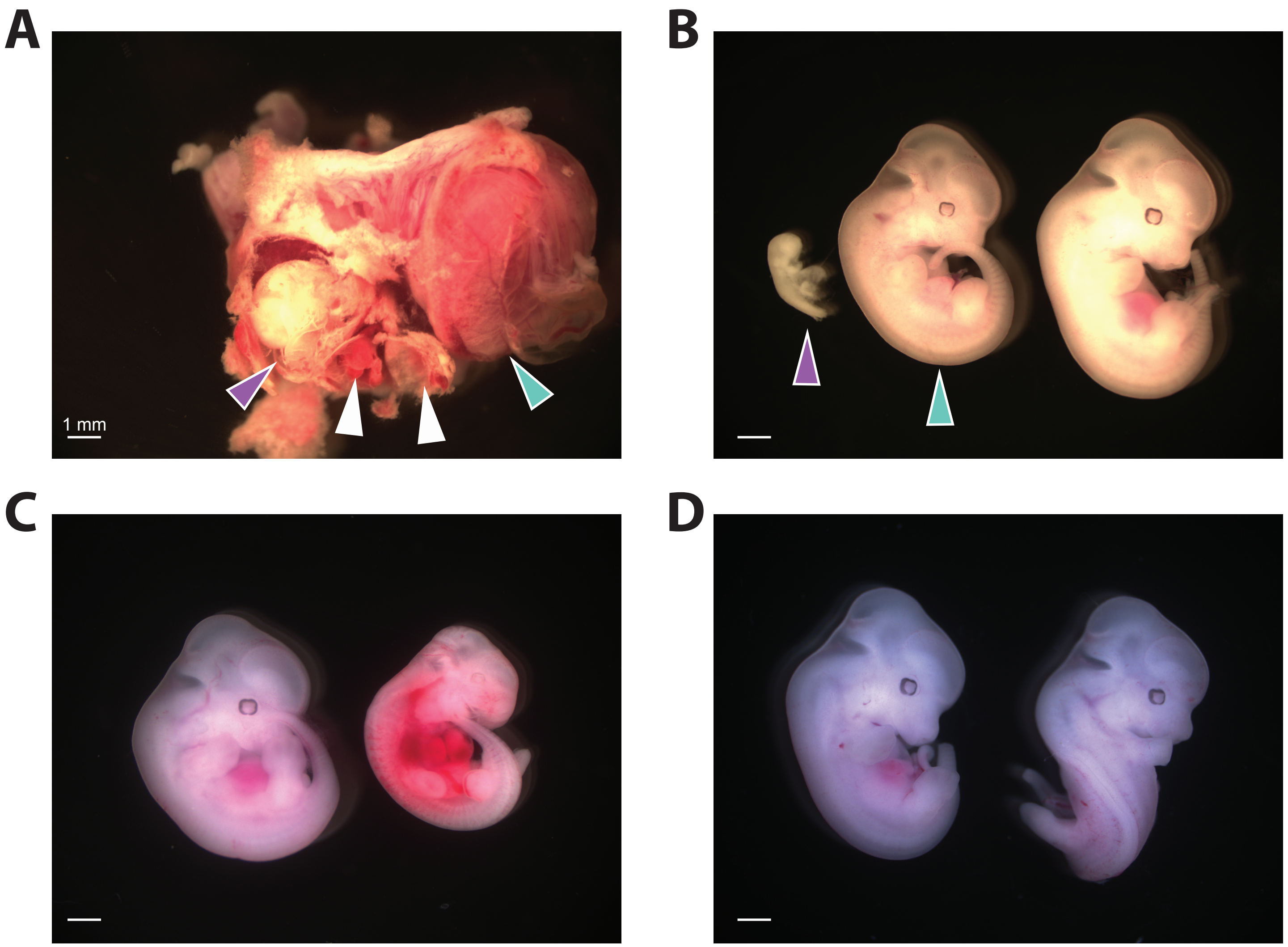


**Supplemental Figure S1:** Range of embryonic phenotypes observed in decidual masses.
A. Dissected implantation sites showing multiple co-implantion events, including two resorbed embryos (white arrows), one severely growth-retarded embryo (purple arrow), and one smaller fetoplacental unit (turquoise arrow) with embryo removed for clarity.
B. Embryos dissected from (A). Colored arrows indicate site of origin from (A). Normal embryo from same pregnancy shown for comparison (far right).
C. A normal (left) and underdeveloped (right) embryo from a close pair.
D. Improper embryo development with twisted body axis (right) associated with mechanical strain during growth with fused placentas.
