## Supplemental Fig. S2 for "Uterine injury during diestrus leads to embryo spacing defects and perturbations in the COX pathway in subsequent pregnancies"

### Supplemental Figure S2

| | | ♂♂ | ♀♀ | ♀♂ |
|---|---|---|---|---|
| 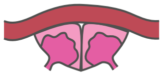 | fused* | 8 | 0 | 9 |
| 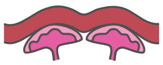 | separate# | 6 | 7 | 6 |
| | total^ | 14 | 7 | 15 |

*$p = 0.02$, $\chi^2$-test
#$p = 0.26$, $\chi^2$-test
^$p = 0.16$, $\chi^2$-test

**Supplemental Figure S2:** Embryo sexes within decidual masses.
Table indicating numbers of fused and separate embryos within decidual masses that were either both males, both females, or of mixed sexes.
