## Supplemental Fig. S3 for "Uterine injury during diestrus leads to embryo spacing defects and perturbations in the COX pathway in subsequent pregnancies"

### Supplemental Figure S3

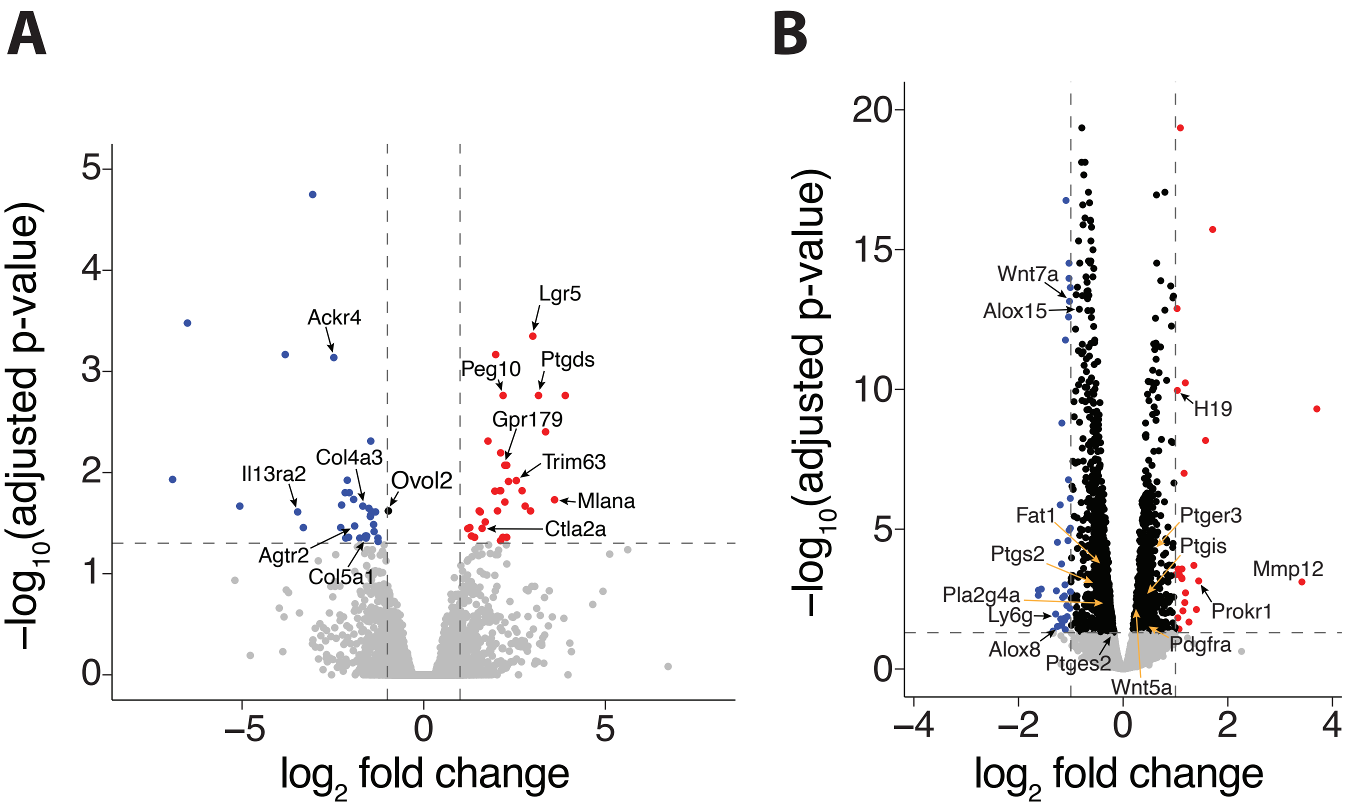


**Supplemental Figure S3:** Differential gene expression analysis.
A. Volcano plot depicting differentially-expressed genes (DEGs) between estrus-injured and -uninjured horns.
B. Volcano plot depicting differentially-expressed genes (DEGs) between diestrus-injured and -uninjured horns.
