## Supplemental Fig. S4 for "Uterine injury during diestrus leads to embryo spacing defects and perturbations in the COX pathway in subsequent pregnancies"

### Supplemental Figure S4

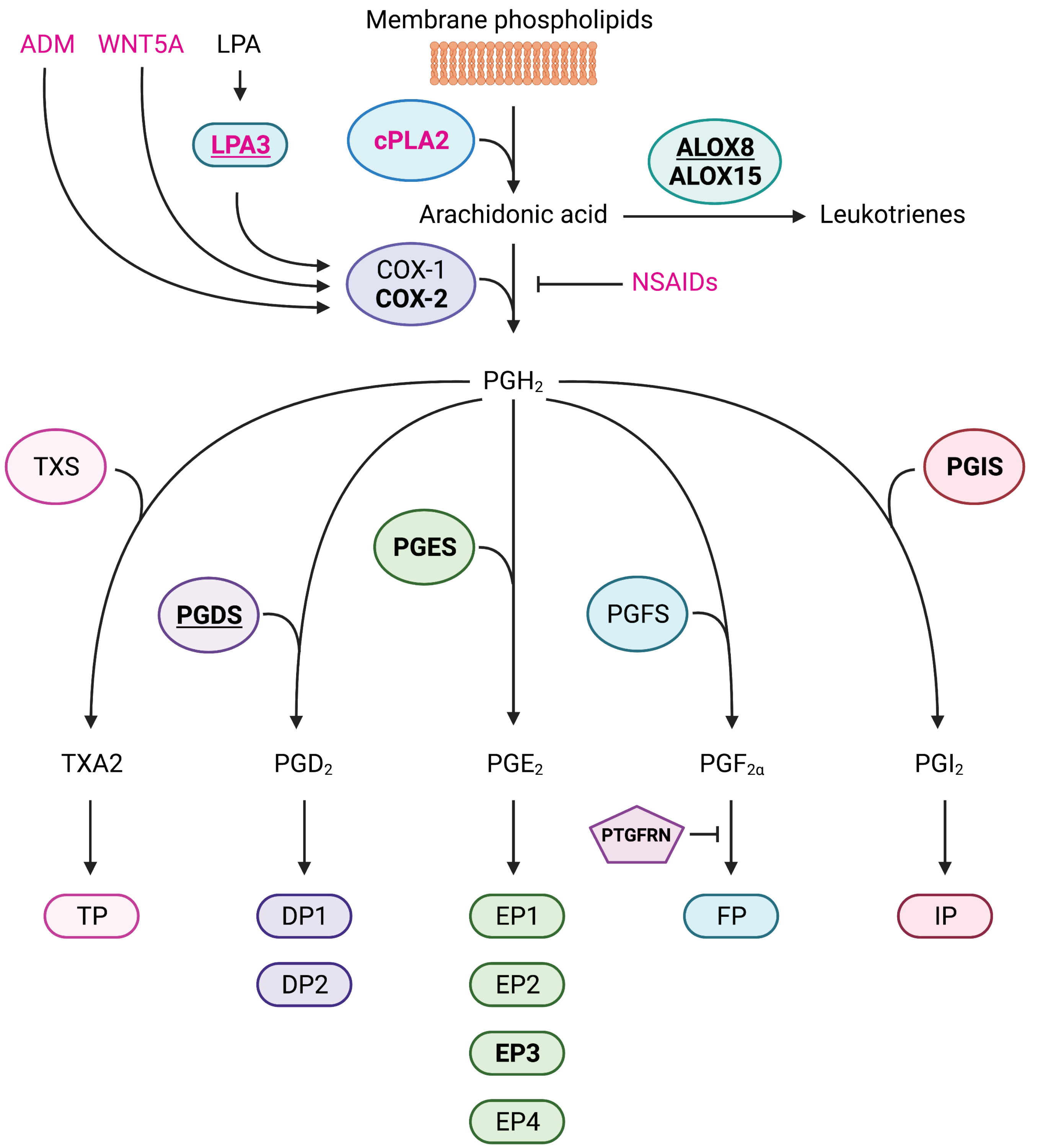


**Supplemental Figure S4:** Diagram of major components of the COX pathway.
Genes dysregulated in diestrus-injured horns are shown in bold, with genes >2-fold change underlined.
Enzymes and factors previously demonstrated to affect embryo spacing are shown in pink text.
Abbreviations: ADM = adrenomedullin, cPLA2α = cytosolic phospholipase A2, COX-2 = cycloooxygenase-2, ALOX8 = arachidonate 8-lipoxygenase, ALOX15 = arachidonate 15-lipoxygenase, TXS = thromboxane-A synthase, PGDS = prostaglandin-D synthase, PGES = prostaglandin-E synthase, PGFS = prostaglandin-F synthase, PGIS = prostaglandin-I synthase, TXA2 = thromboxane A2, $PGD_2$ = prostaglandin $D_2$, $PGE_2$ = prostaglandin $E_2$, $PGF_{2α}$ = prostaglandin $F_{2α}$, $PGI_2$ = prostaglandin $I_2$, TP = thromboxane receptor, DP1 and DP2 = prostaglandin $D_2$ receptors 1 and 2, EP1-4 = = prostaglandin $E_2$ receptors 1 through 4, FP = prostaglandin $F_{2α}$ receptor, IP = prostaglandin $I_2$ receptor, NSAIDs = non-steroidal anti-inflammatory drugs.
